# Gut microbiome shifts with urbanization and potentially facilitates a zoonotic pathogen in a wading bird

**DOI:** 10.1101/718213

**Authors:** Maureen H. Murray, Emily W. Lankau, Anjelika D. Kidd, Catharine N. Welch, Taylor Ellison, Henry C. Adams, Erin K. Lipp, Sonia M. Hernandez

## Abstract

Microbial communities in the gastrointestinal tract influence many aspects of host health, including metabolism and susceptibility to pathogen colonization. These relationships and the environmental and individual factors that drive them are relatively unexplored for free-living wildlife. We quantified the relationships between urban habitat use, diet, and age with microbiome composition and diversity for 82 American white ibises (*Eudocimus albus*) captured along an urban gradient in south Florida and tested whether gut microbial diversity was associated with *Salmonella enterica* prevalence. Shifts in community composition were significantly associated with urban land cover and, to a lesser extent, diets higher in provisioned food. The diversity of genera was negatively associated with community composition associated with urban land cover, positively associated with age class, and negatively associated with *Salmonella* shedding. Our results suggest that shifts in both habitat use and diet for urban birds significantly alter gut microbial composition and diversity in ways that may influence health and pathogen susceptibility as species adapt to urban habitats.

## Introduction

Living in urban areas can have important consequences for wildlife physiology and/or health through changes in habitat use, interactions, and diet [1]. For example, urban birds exhibit higher stress levels and more frequent interactions with non-native species [2]. Similarly, some species experience increased exposure to pollutants and reduced immune function [3]. The provisioning of urban wildlife, either intentionally (e.g., bird feeders) or unintentionally (e.g., garbage), can also promote the use of novel foods [4] and habitat types, such as manicured lawns [5].

Such shifts in wildlife physiology and ecology may, in turn, impact the diverse microbial community inhabiting the host gastrointestinal tract, or gut microbiome. The composition and diversity of the gut microbiome regulates several aspects of host health including the production of vitamins, metabolism, and dietary efficiency (reviewed by [6]). The gut microbiome also mediates host susceptibility to infection and pathogen colonization by initiating proper development of the immune system and the production of antimicrobial metabolites (reviewed by [7]). A healthy, diverse gut microbiome can also act as a barrier to infection by competing, either directly or indirectly, with pathogens for space or resources [8]. This has direct impacts on health. For example, disrupted gut microbial communities are more likely to become colonized by enteric bacteria such as *Clostridium difficile* [9].

The importance of the gut microbiome to host health has led to many studies on humans and livestock attempting to disentangle the relative influence of host age, diet, and environment on microbial communities. These effects can occur through accumulation, whereby older individuals typically have more diverse microbiomes [10]. Host diet can also influence microbiome composition and diversity through macronutrient balance [11]. For example, diets higher in protein are typically associated with certain phyla and increased diversity [12]. Lastly, hosts are in constant contact with microorganisms in their environments from various substrates, which can shape the acquisition and composition of intestinal microbiota (e.g., [13,14]).

The roles of environmental exposure and diet in shaping the host microbiome may cause the gut microbiome to act as an important mediator for how urban environments or changing diets may affect host health. The shifts in diet and habitat exhibited by wildlife in urban environments likely influence their gut microbial communities as they do for animals in captivity [15–18] and humans in urban vs. rural communities [19]. While a rapidly growing body of literature has characterized the avian microbiome for many species [20], little is known about how the novel foods and habitat types available to urban birds may shape microbiome diversity and composition. For example, urban house sparrows have less diverse microbiomes than their rural counterparts due to several potential mechanisms [21].

Such shifts in diet and habitat use are apparent for the American white ibis (*Eudocimus albus*), a mid-sized carnivorous wading bird native to the southeastern United States. Up until the 1960’s, Florida was the primary breeding site for ibis on the mainland USA, but following dramatic wetland loss and development, breeding colonies experienced a 90% decline of nesting pairs in Florida. In the last 20 years, ibis have been observed foraging in urban environments in cities across south Florida [22,23]. Our previous work in this system has demonstrated that ibises inhabiting more developed urban areas are more likely to consume provisioned food, such as bread, but have poorer body condition [24] and are more likely to shed *Salmonella enterica* [23]. Almost half of urban ibis *Salmonella* genotypes matched those for human salmonellosis cases, indicating ibis have the ability to act as reservoirs of salmonellae for people, or share a common environmental source [23]. In a more robust comparison of *Salmonella* shedding by ibis from developed and natural environments, site fidelity to contaminated urban parks was a significant predictor for *Salmonella* prevalence [25]. These observed changes in the diets and health of ibis with urbanization may be indicative of concurrent changes in the composition or diversity of the gut microbiome for urban birds.

In this study, we characterized the gut microbiome of white ibises with the goal of determining how gut microbiome community composition and alpha diversity changed with urbanization, the consumption of provisioned food, age, and sex. We then tested whether any differences in composition or diversity were associated with the prevalence of *Salmonella* spp. To do so, we sampled white ibises across an urban gradient in South Florida and collected feces for *Salmonella* isolation and microbiome analyses and blood for stable isotope analysis of diet.

## Materials and Methods

### Ibis capture and sample collection

We captured ibises in Palm Beach and adjacent counties in South Florida between October 2015 and March 2017 (Fig 1). South Florida has a tropical climate with a high biodiversity of wading birds and annual fluctuations in water levels between the dry (November – April) and rainy (May – October) seasons. Palm Beach County has dense urban developments along the coast progressing to agricultural development and wetlands further inland. We captured ibises at 15 sites where we had observed foraging ibises and that represented an urban land cover gradient (0-93% urban land cover, see [24] for details on land cover analysis). These sites included five restored or constructed wetlands, two wildlife rehabilitation centers, a landfill, a zoo, and six urban parks. We captured ibises in wetlands using mist nets with decoys and at more urban sites we used baited manually-operated flip traps and nylon slip-knot leg lassos [24].

**Figure 1.**
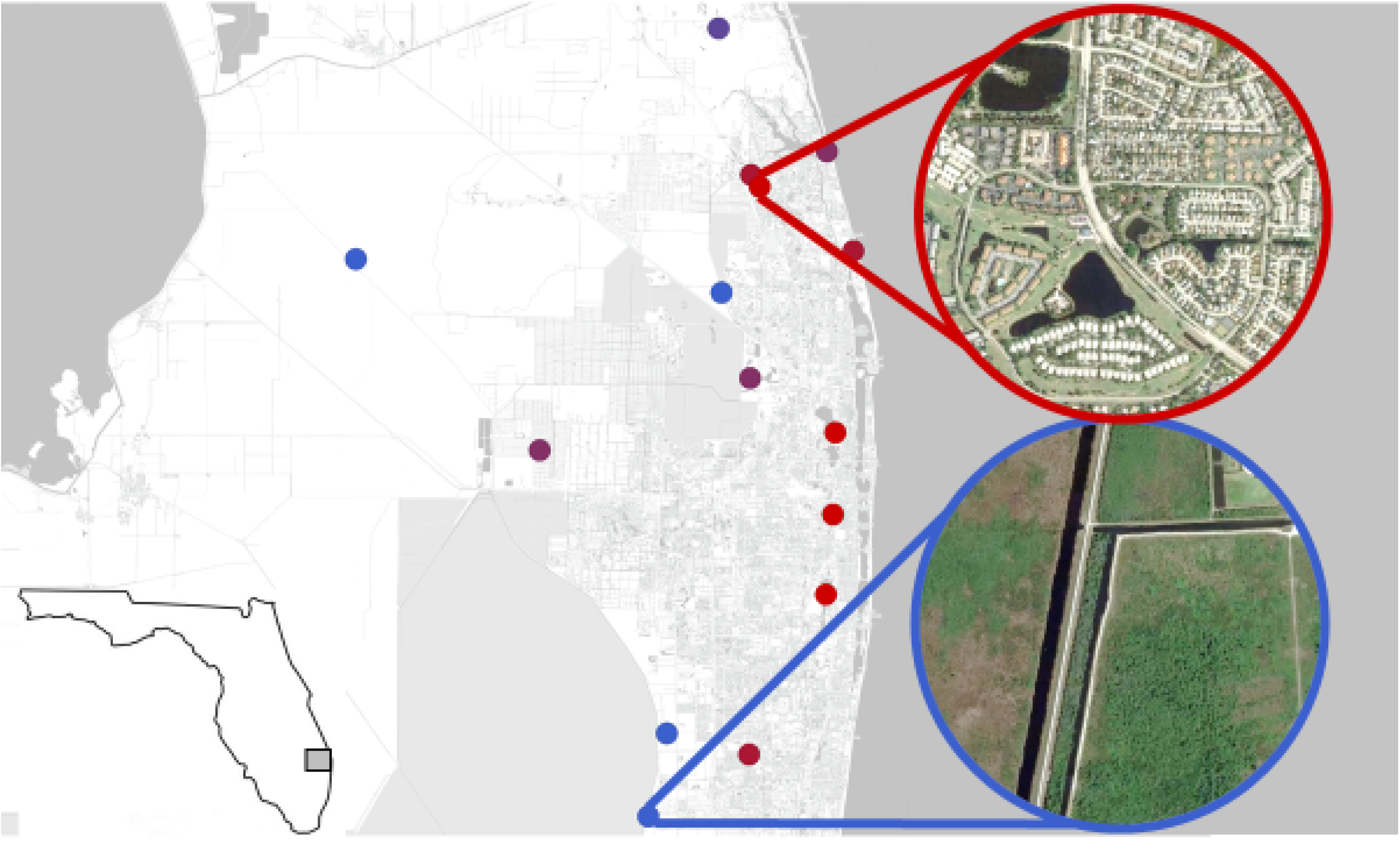
Map of 15 sampling sites on the eastern coast of South Florida where American white ibises were captured to collect fecal specimens to determine their gastrointestinal microbiome. Sampling sites represented a gradient of urban land cover from 0% (blue) to 91% (red; see also Table S1). Inset circles demonstrate the range of difference in urban habitat across the urban land cover gradient, ranging from open wetland (blue inset) to densely inhabited residential areas (red inset).

We assessed ibis age using plumage as adults (≥3 years, all white feathers) or subadults (some brown feathers) [26] and recorded mass, culmen length, wing chord length, tarsus length and width to calculate body condition. We then collected ≤1% body weight of blood from each bird from the jugular or metatarsal vein. We used whole blood samples to confirm ibis sex using standard molecular techniques [27] and for stable isotope analysis of diet (see below). If a captured ibis defecated onto a clean surface, we collected ≥0.25 g of feces (0.63 ± 0.05, mean ± S.E.) using a sterile cotton tipped applicator for microbiome analysis. We also collected approximately one gram of fresh feces into 10 ml of dulcitol selenite broth for *Salmonella* isolation.

### Molecular characterization of bacterial and fungal communities

#### Sample processing and extraction

Fecal samples for gut microbiome analyses were stored in microcentrifuge tubes at −20C in the field and subsequently at −80C until DNA extraction. Up to 0.25 mg of sample was aliquoted into 96 well extraction plates for submission to the University of Wisconsin Biotechnology Center for extraction and sequencing. Bacterial and fungal DNA were extracted using the PowerMag microbial DNA isolation kit following the manufacturer’s protocol (Mo Bio Laboratories, Inc.). Extracted DNA was cleaned using a SpeedBead Clean-up protocol (GE SeraMag Beads, GE Healthcare, Chicago, IL) prior to library preparation.

#### Construction and sequencing of v3-v4 16s metagenomic libraries

DNA concentration was verified fluorometrically using either the Qubit^®^ dsDNA HS Assay Kit or Quant-iT^™^ PicoGreen^®^ dsDNA Assay Kit (ThermoFisher Scientific, Waltham, MA, USA). Samples were prepared in a similar process to the one described in Illumina’s 16s Metagenomic Sequencing Library Preparation Protocol, Part # 15044223 Rev. B (Illumina Inc., San Diego, California, USA) with the following modifications: The 16S rRNA gene V3/V4 variable region was amplified with fusion primers (forward primer 341f: 5’-ACACTCTTTCCCTACACGACGCTCTTCCGATCT(N)3/6CCTACGGGNGGCWGCAG-3’, reverse primer 805r: 5’-GTGACTGGAGTTCAGACGTGTGCTCTTCCGATCT(N)3/6GACTACHVGGGTATCTAAT CC-3’). Region specific primers were previously described in [28] (underlined sequences above), and were modified to add 3-6 random nucleotides ((N)3/6) and Illumina adapter overhang nucleotide sequences 5’ of the gene-specific sequences. Following initial amplification, reactions were cleaned using a 0.7x volume of AxyPrep Mag PCR clean-up beads (Axygen Biosciences, Union City, CA). In a subsequent PCR, Illumina dual indexes and Sequencing adapters were added using the following primers (Forward primer: 5’-AATGATACGGCGACCACCGAGATCTACAC[55555555]ACACTCTTTCCCTACACGACG CTCTTCCGATCT-3’, Reverse Primer: 5’-CAAGCAGAAGACGGCATACGAGAT[77777777]GTGACTGGAGTTCAGACGTGTGCTC TTCCGATCT-3’, where bracketed sequences are equivalent to the Illumina Dual Index adapters D501-D508 and D701-D712,N716,N718-N724,N726-N729). Following PCR, reactions were cleaned using a 0.7x volume of AxyPrep Mag PCR clean-up beads (Axygen Biosciences). Quality and quantity of the finished libraries were assessed using an Agilent DNA 1000 kit (Agilent Technologies, Santa Clara, CA) and Qubit^®^ dsDNA HS Assay Kit (ThermoFisher Scientific), respectively. Libraries were pooled in an equimolar fashion and appropriately diluted prior to sequencing. Paired end, 300 bp sequencing was performed using the Illumina MiSeq Sequencer and a MiSeq 600 bp (v3) sequencing cartridge. Images were analyzed using the standard Illumina Pipeline, version 1.8.2. OTU assignments and diversity plots were created using QIIME analysis pipeline [29].

#### Bioinformatics

Output sequencing files provided by the University of Wisconsin Biotechnology Center were separated by barcode sequences by the Biotech center, resulting in 192 pairs of forward and reverse read fastq files. Paired sequences were joined, quality filtered, clustered into operational taxonomic units, assigned to taxonomic identities, and aligned to create a phylogenetic tree using the QIIME pipeline [29]. Paired end reads were joined using the multiple_join_paired.py script and default parameters, separately for the two batches of files coming from the two Miseq runs. Unpaired sequences were discarded. Joined consensus sequences were quality filtered and combined into one fasta file using the multiple_split_libraries_fastq.py script with the “sampleid_by_file” option, again separately for the two Miseq runs, using default parameters. The two resulting combined files were then concatenated into one final file. Because we used the same sample identifiers on the two separate Miseq runs, by concatenating the output files from the two split_libraries script we effectively pool the sequences from both Miseq runs for each sample.

We clustered the resulting sequences into operational taxonomic units with 97% identity using the parallel_pick_open_reference_otus.py script with the UCLUST method. Open reference OTU clustering works by first comparing each of our query sequences against a reference database – we used the Greengenes 97% identity database [30]. Then, any sequences not matching to an entry in the reference database at 97% identity are separately clustered first by creating new reference databases from a random draw of the sequences for two iterations (pulling 0.1% of the unclustered sequences each time). Finally, any sequences not assigned to an OTU after these steps are clustered using de novo clustering. This process was run on 12 nodes on the University of Wisconsin Center for High Throughput Computing.

OTUs were assigned to the highest confident taxonomic resolution using the RDP Naïve Bayesian Classifier [31], using the assign_taxonomy.py script in QIIME. A final community matrix (samples as rows, OTUs as columns, with the number of sequences per OTU detected in each sample as cells) was created using the make_otu_table.py script.

Representative sequences for each OTU were aligned using the align_seqs.py script, using the PyNAST method using default parameters. Aligned sequences were used to create a phylogenetic tree, using the make_phylogeny.py script with the FastTree method [32]. For phylogenetically informed analyses, the OTU table was filtered to remove any OTU’s whose representative sequences could not be aligned. The reshape2 package in R [33] was used to collapse OTUs into higher taxonomic levels for analysis (see Table S1 and Figure S1 for a summary of bioinformatics output at the genus level, which was used for analysis).

### Salmonella isolation

We isolated *Salmonella* from fecal samples by modifying previously published protocols [34]. Briefly, 1 g of feces was placed in dulcitol selenite pre-enrichment broth and was maintained at ambient temperature for approximately 2-4 days until they were shipped to the University of Georgia (Athens, Georgia) for culture. We then transferred 100μl of turbid pre-enrichment broth to 10 ml of Rappaport-Vassiliadis (RV) broth and incubated for ~24 h at 43 °C (Oxoid, Hampshire, UK). We streaked for isolation from each RV enrichment onto xylose lysine tergitol 4 (XLT-4) agar plates and incubated for 24 ± 2 h. Colonies suspected to be *Salmonella* based on morphology (black colonies or yellow colonies with a black center) streaked again for isolation to obtain a pure colony. Each isolate was then confirmed by characteristic growth on CHROMagar© Salmonella-Plus after 24 h of incubation at 37 °C [34] (CHROMagar, Paris, France). We confirmed a subset of isolates as *Salmonella* via PCR [25,34]. Confirmed *Salmonella* isolates were stored at −80 °C.

### Stable isotope analysis

We quantified the proportion of provisioned food (i.e., bread and chips) in ibis diet using δ^13^C and δ^15^N stable isotope analysis of red blood cells. These isotopes reflect the carbon sources and trophic level of consumer diets; δ ^13^C differs for marine and terrestrial communities and C4 plants such as corn in processed foods [35], and δ^15^N is biomagnified in predators [36]. Avian red blood cells have a turnover rate of approximately two months [37] and so provide an estimate of assimilated diet over that period. We estimated the importance of provisioned food, freshwater and terrestrial invertebrates, marine invertebrates, landfill refuse, and fish by creating five-source mixing models using the R package SIAR [38]. Samples were analyzed following standard methods at the Center for Applied Isotope Studies at the University of Georgia (see [24] for in-depth description of diet sources and laboratory methods).

### Statistical analyses

Microbiome data were analyzed at the genus and phyla levels. Microbial community composition was analyzed at the site level (i.e., relative abundance of each taxon was averaged across individual samples within a site) to control for pseudoreplication when comparing microbiome characteristics to site-level characteristics. Similarity among sites in average genus-level microbiome composition was visualized using principal coordinates analysis using the capscale function in the vegan package in R [39]. Associations between the microbiome and the geographic and demographic explanatory variables were assessed using environmental fits performed with the envfit function in the vegan package in R.

At the phylum level, association between the site-average relative abundances of each phylum and landscape urbanization were visualized using the heatmap function in the pheatmap package in R [40]. Significance of the associations between site-average relative abundance of each phylum and landscape urbanization was tested using a sample-size weighted correlation using the wtd.cor function in the weights package in R [41] with adjustment of significance thresholds using false discovery rates [42] to control for multiple comparisons. Only phyla with an average relative abundance ≥0.1% were included in this analysis.

The relationships among microbial community characteristics at each site (e.g., Shannon diversity at the genus level and the ordination axes as univariate measures of community composition) and key environmental variables (landscape urbanization, dietary provisioning, and *Salmonella* prevalence) were assessed using linear or logistic regression as appropriate based on exploratory graphical analysis of the pair-wise relationships. Linear regression was performed using the lm function the stats package in R [43]. Linear regressions were performed on site-level measures and were weighted by samples size at each site. Logistic regression was performed on the individual sample characteristics for genus-level Shannon diversity and *Salmonella* presence using the glm function in the stats package in R with a binomial distribution. This relationship was visualized at the site level to facilitate comparison with other site-level correlations.

Explanatory variables were correlated to both microbiome measures and each other, making it difficult to determine which variables linked directly to patterns in microbial community measures and *Salmonella* prevalence. To address this issue, we performed structural equation modeling to better understand the relationships among the variables. Structural equation modeling was performed using the lavaan [44] and lavaan.survey packages [45]. We identified the best fit model starting from a complete model with all hypothesized relationships and interactions and then performed a manual backwards selection removing non-significant relationships from the model until the model fit stabilized with acceptable model fit parameters. This final model was tested further by removing one or more relationships from the model and comparing the model fit parameters for decline or improvement in model stability.

## Results

We sampled 82 white ibises at 15 sites (5.3 ± 4.0 birds per site) and sampled more adult females than other age and sex classes (adult females = 36, adult males = 20, subadult females = 17, subadult males = 9). Microbiome sequence analysis of white ibis feces yielded on average 104,695 sequence reads (± 6077 SEM) and 351.6 genera (± 19 SEM) per sample (Figure S1, Table S2).

### Changes in composition

The dominant shift in gut microbiome composition at the genus level correlated to the percent of urban landcover, the average percentage of diet from provisioned food, and average diversity (Shannon) at the genus level (Figure 2: Environmental Fit – Percent urban landcover p<0.0001, R^2^=0.835; Average percent of diet provisioned p=0.021, R^2^=0.493; Percent of samples from adults p=0.103, R^2^=0.314; Shannon diversity of genera p=0.014, R^2^=0.519). Urban landcover and the average percentage of diet from provisioned foods in turn were moderately correlated (sample-size adjusted R^2^=0.304, p=0.020).

**Figure 2.**
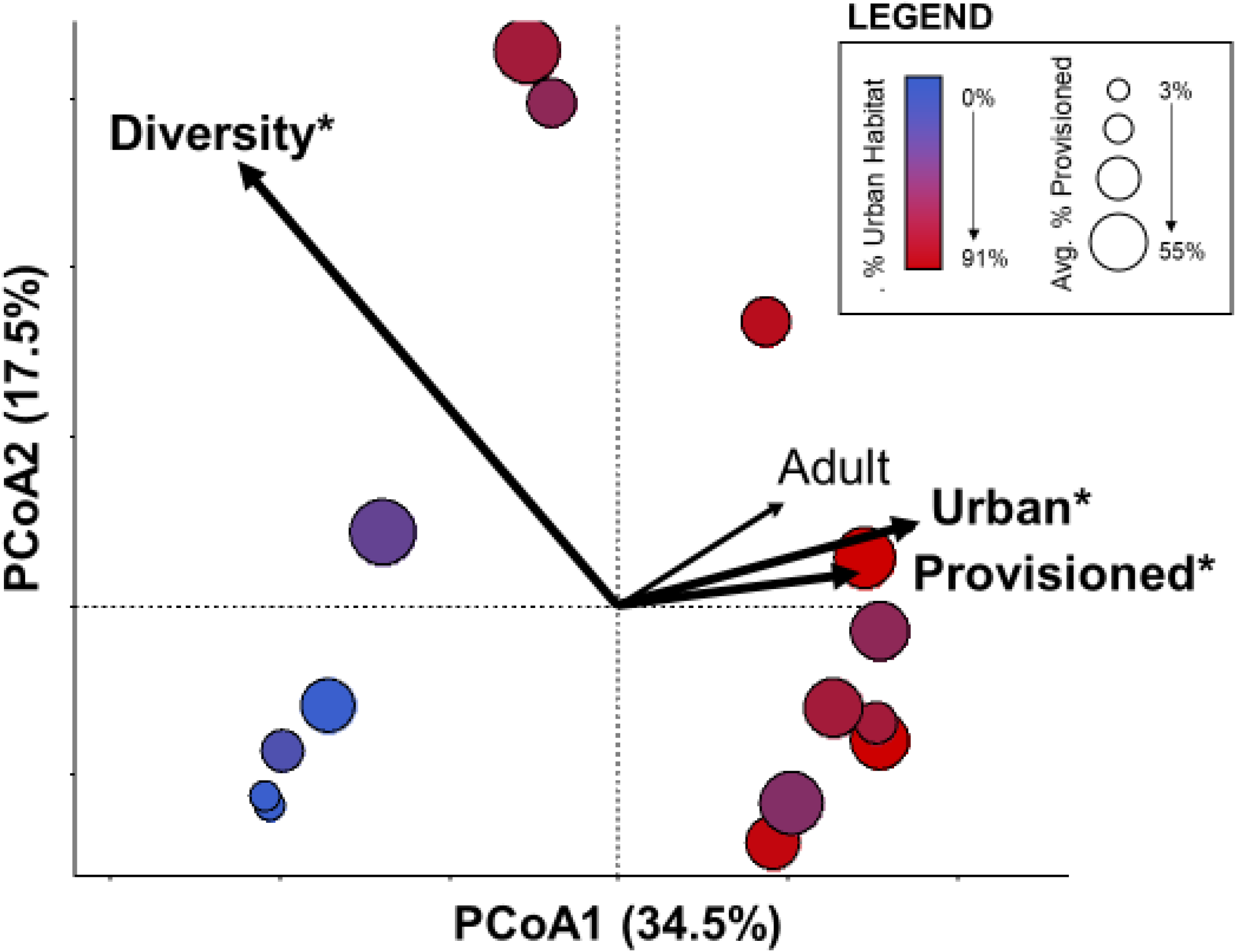
Relationship between bacterial community composition to urbanization, dietary provisioning, age, and genus diversity. Principal coordinates analysis demonstrated site-level patterning in bacterial community composition at the genus level that corresponded to urban land cover (red to blue gradient) and use of provisioned food by ibises (circle size) at capture sites, but not age composition of sampled birds (See text for statistics). Genus-level Shannon diversity associated with the community composition, increasing in the opposite direction of these independent variables.

Urban land cover and dietary provisioning were primarily associated with changes in genus-level bacterial community composition along the first ordination axis (PCoA1) (Figure 2, R^2^ = 0.82, F = 59.0, p < 0.001). This shift in composition corresponded to notable changes in the relative abundance of several bacterial phyla (Figure 3, Table S3). The most abundant phyla in white ibis feces were Firmicutes (mean percentage of sequences across all samples: 32.5%), Proteobacteria (22.6%), and TM7 (22.6%); however, average relative abundance of dominant phyla varied considerably across the urbanization gradient (Figure 3, Table S3). Firmicutes and Cyanobacteria significantly decreased in relative abundance while Proteobacteria, TM7, Bacteroidetes, OP11, and TM6 increased in relative abundance with increasing urban land cover (Table S3).

**Figure 3.**
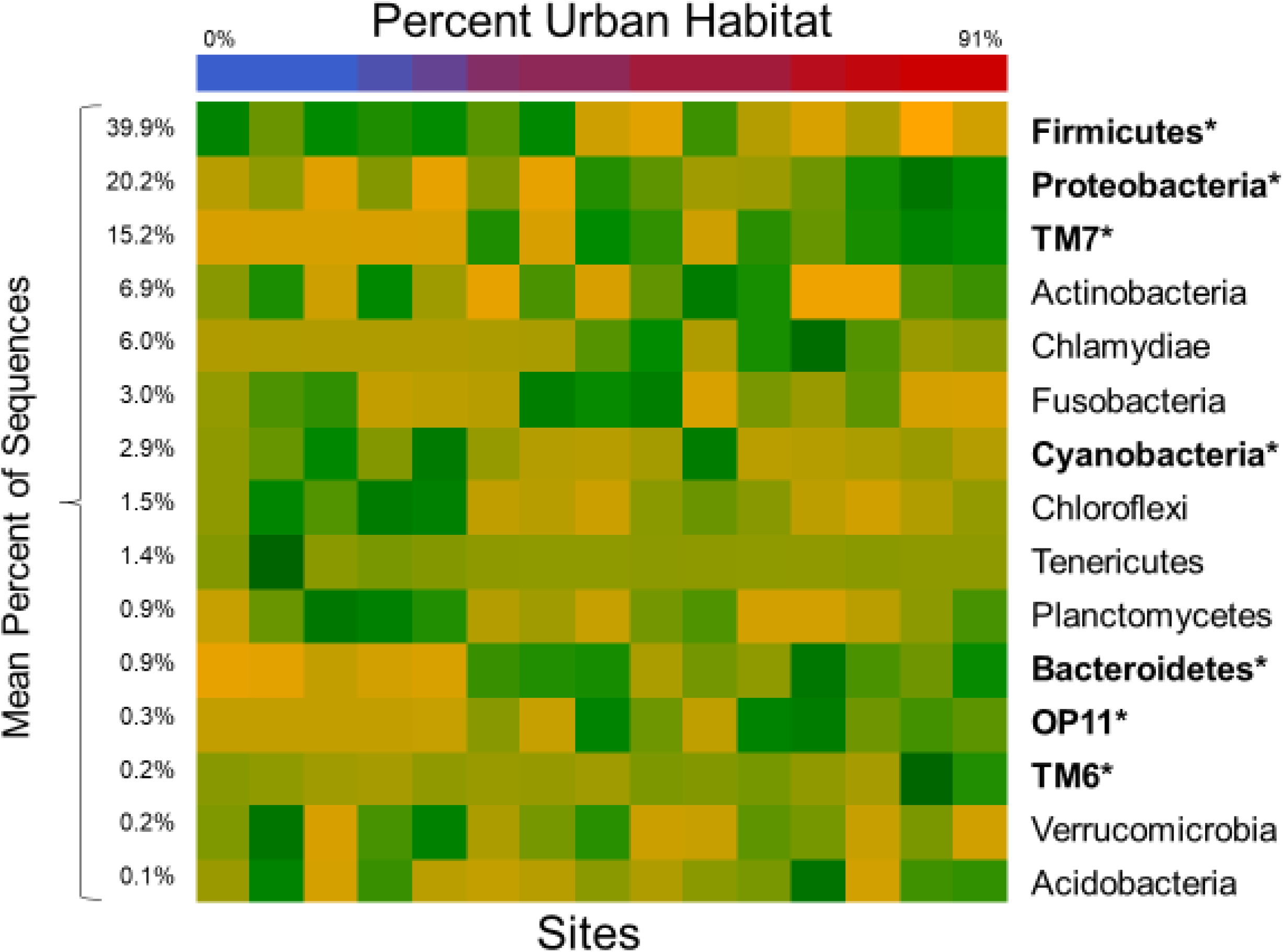
Correlation between urban landcover and average relative abundance of dominant bacterial phyla. The heat map shows several bacterial phyla (bolded) significantly increased (green) or decreased (yellow) in relative abundance across the urban landcover gradient (top bar, on the same color scale as Figure 2).

### Changes in diversity

Unlike bacterial composition, changes in bacterial genus diversity were associated with individual ibis biology rather than capture site. These changes in genus diversity, as measured by the Shannon diversity index, were not associated with urban land cover or dietary provisioning (Urban land cover: R^2^ = 0.04, F_1,13_ = 1.16, p = 0.30, Figure 4a; Provisioning: R^2^ = 0.03, F_1,13_ = 0.81, p = 0.38, Figure 4b). Rather, diversity was significantly correlated with microbiome community composition (PCoA1) (R^2^ = −0.29, F_1,13_ = 5.2, p = 0.04; Figure 4c). As predicted, ibis age, measured as the proportion of adult ibis captured at a particular site, was positively correlated with genus diversity (R^2^ = 0.16, p < 0.001). Mean bacterial diversity was negatively correlated with *Salmonella* prevalence (Figure 4d), such that ibises with lower genus diversity were more likely to be shedding *Salmonella* (β = −2.10 ± 0.99, z = −2.12, p = 0.03).

**Figure 4.**
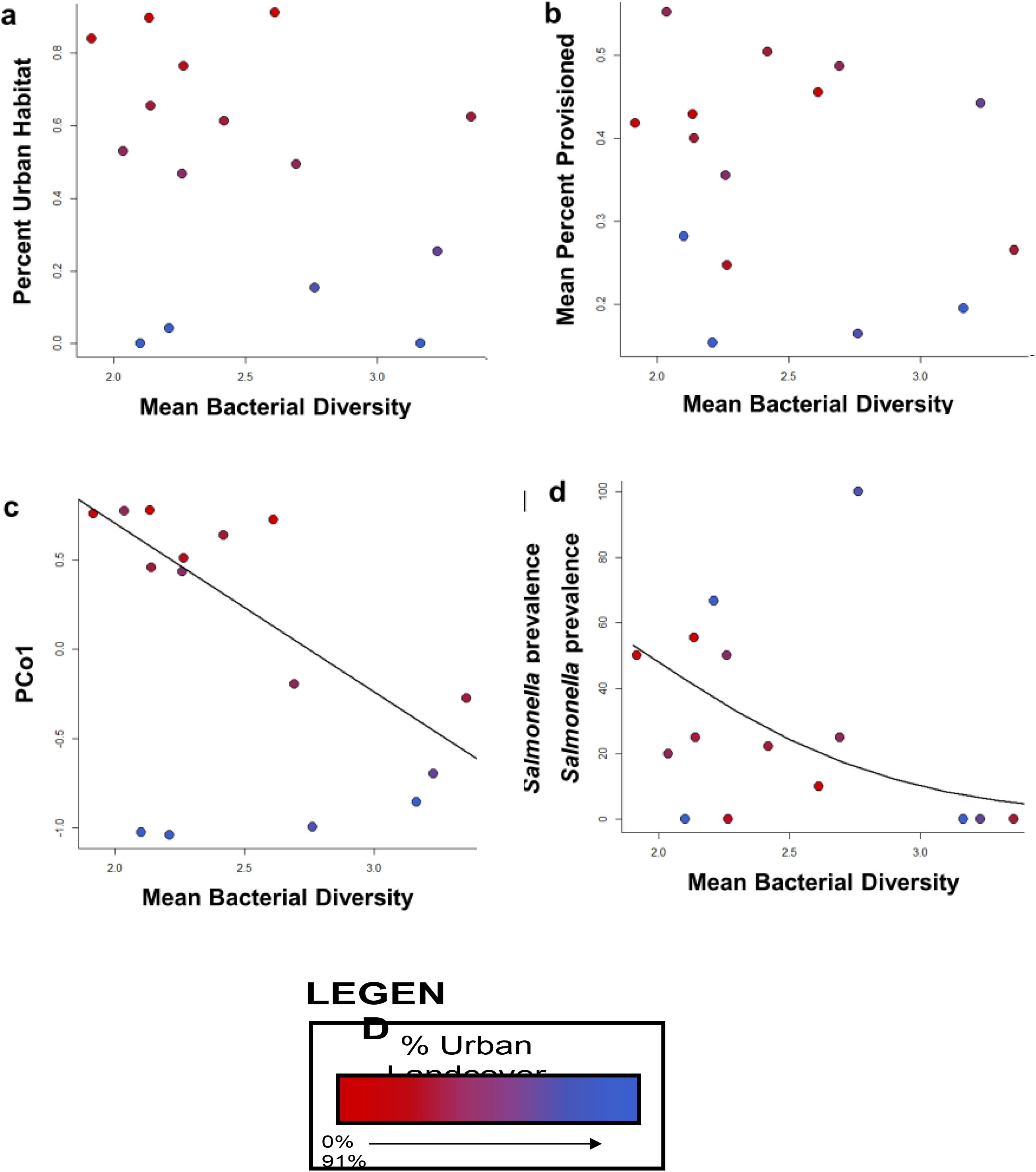
Correlation of bacterial diversity to urbanization (a), dietary provisioning (b), and bacterial community composition (c), and *Salmonella* prevalence (d). Circles represent average values for 15 capture sites and colored along an urban gradient from red (most urban) to blue (least urban). Bacterial diversity was not significantly correlated to landscape and dietary factors but was significantly associated with overall community composition and *Salmonella* prevalence (see text for statistics).

### Diversity and *Salmonella* prevalence

Given the co-linearity between several of our explanatory variables and community composition, we developed a structural equation model to better understand how these variables linked through gut community composition to associations with *Salmonella* prevalence (Structural equation model weighted by sample size at each site: Robust Chi-squared goodness-of-fit p=0.466; Comparative Fit Index =1.000; Tucker-Lewis Index=1.032, Fig 5). The best model indicated that bacterial community composition was associated with both the degree of urbanization surrounding the sampling site and to a lesser extent the percentage of the diet attributed to provisioned food, where urbanization acted both directly on composition and indirectly through effects on diet (Urbanization: β = 1.18 ± 0.21, z = 5.64, p < 0.001, Diet: β = 1.51 ± 0.69, z = 2.18, p = 0.029, Figure 5). Bacterial community composition (PCo1) was negatively associated with average genus-level diversity, and diversity was positively associated with the proportion of adult ibis at each site (PCoA1: β −0.39 ± 0.15, z = −2.62, p = 0.009, Proportion adult: β = 0.65 ± 0.14, z = 4.68, p < 0.001, Figure 5). Genus-level diversity was then negatively associated with *Salmonella* prevalence (β = −0.41 ± 0.17, z = −2.42, p=0.016; Figure 5).

**Figure 5.**
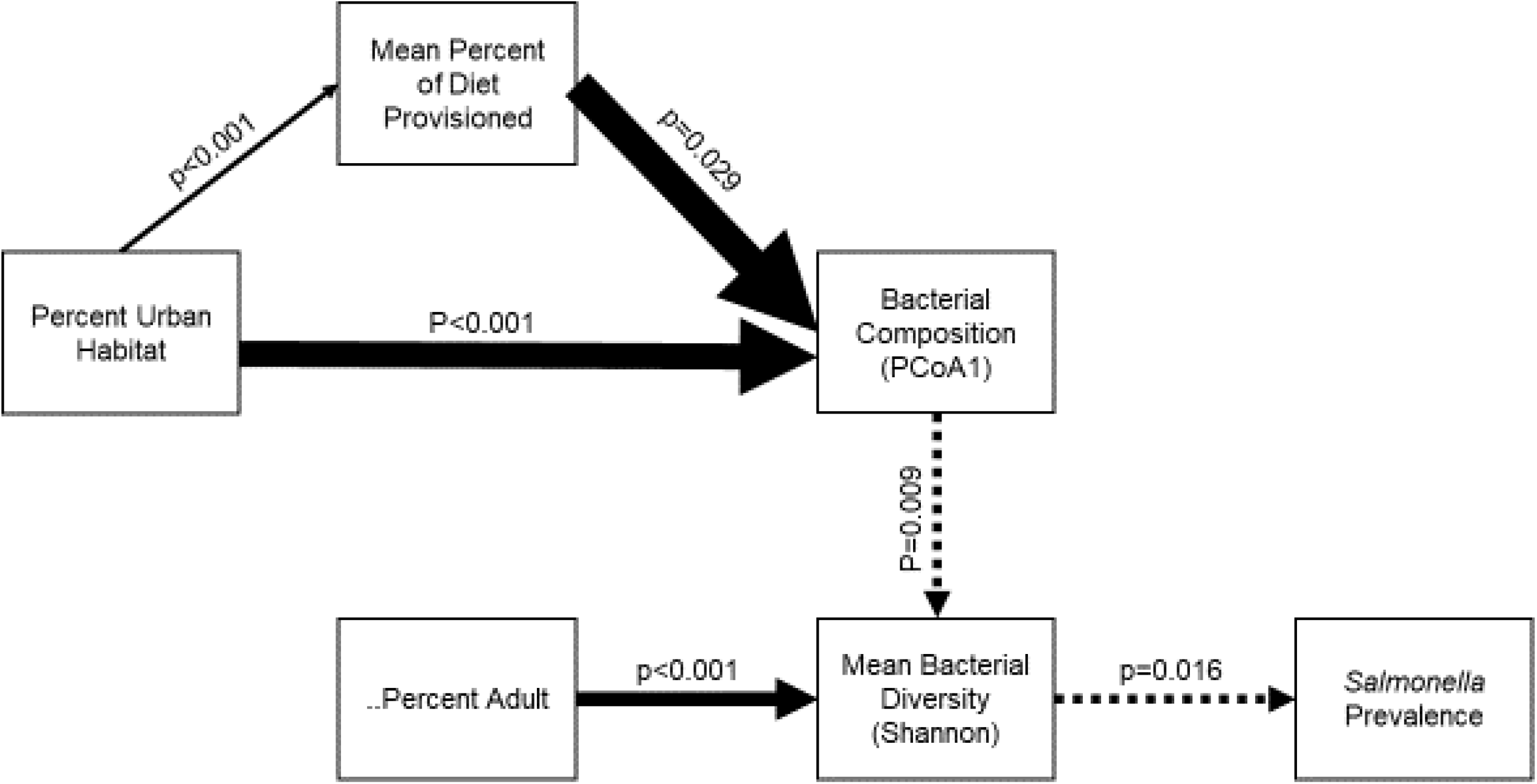
Path analysis of factors associated with *Salmonella* prevalence in American white ibises. Path analysis showing significant positive (solid arrows) and negative relationships (dashed arrows) between urban land cover, ibis use of provisioned food, and ibis age with microbial composition (ordination axis values), diversity (Shannon index of genus diversity), and Salmonella prevalence. Thicker arrows indicate larger effect sizes.

## Discussion

In this study, we examined changes in the composition and diversity of the gut microbiome with urbanization and food provisioning for the American white ibis, a recently urbanized wading bird. We also tested if these changes were associated with an increased likelihood of shedding *Salmonella*, an opportunistic enteric zoonotic pathogen. Microbiome community composition was strongly associated with urban land cover and, to a lesser extent, the assimilation of provisioned food, such as bread. Microbial diversity was negatively associated with these changes in community composition, and ibises with less diverse microbiomes were more likely to shed *Salmonella*.

Urban land cover was the strongest driver of microbial community composition and is itself associated with many biotic and abiotic factors. Urban ibis in Australia also consume provisioned bread in parks and select for high-carbohydrate rather than high-protein foods [46]. The relative importance of macronutrients is an important predictor of microbiome composition across species [47] and so may be at least partially driving the shifts we observed in phylum relative abundance. Ibises in urban parks may also be exposed to different environmental bacteria when foraging on lawns, managed ponds, and paved surfaces rather than shallow wetlands. Similarly, ibises and other birds in urban areas often interact with non-native or invasive species [48], further increasing their exposure to novel microbiota. Other differences in the behavior or physiology of urban birds, such as increased sedentariness [49] or altered stress levels [2] may alter microbiome composition through habitat exposure and physiological interactions. Regardless of mechanism, the taxonomic shifts we observed in microbiome composition had important consequences for community diversity.

Similarly to community composition, changes in gut microbial diversity can be caused by many different factors relating to bacterial exposure and within-host dynamics. Although urbanization was not directly associated with changes in bacterial diversity, these factors were linked indirectly through changes in composition. These findings are consistent the only other studies examining the shifts in wildlife microbiomes with urbanization, where urban house sparrows (*Passer domesticus*) and urban gulls (*Larus argentatus*) also had less diverse gut microbiomes, potentially due to less varied diets or habitat use [21,50]. Ibises that use less urban areas may be exposed to more diverse environmental bacteria because they exhibit more nomadic movements and use varied habitats such as freshwater and estuarine wetlands, coastal areas, and agricultural areas [25]. Similarly, older individuals of many species, including humans, have more diverse gut microbial communities, likely because of accumulation from environmental exposure over time [6,10].

Perhaps most importantly, lower average microbial diversity was significantly associated with higher *Salmonella* prevalence. In humans and other mammals, gut microbiome perturbations have been associated with an increase in infections and various inflammatory and chronic diseases [17,51–53]. The protective role of microbiome diversity has been demonstrated in many model species. In mice, following experimental treatment with antimicrobials to dramatically alter the composition of the gut microbiome, *Salmonella enterica* serovar Typhimurium or mice experimentally infected with *Clostridium difficile* became “supershedders”, facilitating rapid transmission [54,55]. This link between microbiome diversity and infection has also been demonstrated in other systems—Avian Influenza (AI)-positive mallards had lower OTU diversity than AI-negative birds [56]. The relationship between gastrointestinal microbiome and pathogens in free-living animals is, however, understudied. When the gut microbiome of captive cheetahs was compared to free-living cheetahs in the same habitat, captive individuals had higher abundance of pathogenic bacteria OTUs and modified disease-associated functional pathways [17]. In fact, manipulations of the gut microbiome may offer unique treatment modalities and this has been examined in production animals. For example the administration of prebiotic attenuated *Salmonella* strain or prebiotic galacto-oligosaccharides (GOS) can promote resistance to *Salmonella* colonization via increases of beneficial microorganisms in chickens [57] and similarly, *Lactobacillus* is used as a probiotic to protect against common pathogens like *Salmonella* and *Campylobacter* [58]. And specifically because antimicrobial resistance of *Salmonella* has become an urgent problem, studies on the protection against *Salmonella* colonization and infection through microbiome manipulation chickens abound [59,60]. Yet, studies where the relationships between diet, microbiome and pathogens are examined are extremely limited for free-living populations. Consistent with our results, Jungle crows (*Corvus macrorhyncos*), which mostly consume anthropogenic food, were found to have a relatively high prevalence of pathogenic organisms in their gut microbiome [61].

Relevant to our work, the chicken gastrointestinal microbiome both interacts with the immune system within the lumen of the intestinal tract in various ways, and directly competes with pathogens. For example, greater number of intestinal lymphocytes are present in conventional versus germfree chickens and at the molecular level, the complexity of the T-cell receptor repertoire is influenced by gut microbial diversity. Additionally, in chickens, used both as human animal models and for their own economic impact, pathogen exclusion through competition appears to be the mechanism responsible for lower *Salmonella* colonization rates in older chicks with complex microbiomes [62].

The ability of microbiota to protect the host via resisting the colonization of pathogen parallels the increased resistance to invasion by more diverse ecological communities more generally. There are many studies demonstrating increased invasion success by pathogenic bacteria following a loss of microbial diversity parallels ecological theory that more diverse communities are more resistant to invasion [63]. Similar to other ecological systems, microbial symbionts can protect hosts inadvertently though intense competition for resources from the host’s diet [64]. Likewise, just as invasive species may exploit disturbed communities [65], opportunistic pathogens such as *Salmonella enterica* exploit and more easily colonize disturbed microbial communities [63,66].

Although more diverse microbial communities can protect the host from pathogen colonization, the negative relationship we observed between microbiome diversity and *Salmonella* prevalence may also be caused by microbiome disruption following *Salmonella* colonization. For example, S. Typhimurium can cause dysbiosis by inducing inflammation through the expression of virulence factors [66,67]. While we cannot determine the direction of causation between *Salmonella* prevalence and microbiome diversity with the data we collected in this study, it is a rich avenue for further research in free-living wildlife populations. For example, further characterization of the endogenous microbial communities of different wildlife species can help identify dysbiosis and any threats to conservation or public health [20].

Many studies have demonstrated changes in the prevalence of pathogen prevalence, immune function, and stress in urban wildlife [1,68,69]. Shifts in microbiome diversity with habitat use and diet may be an underappreciated mechanism by which urbanization can affect wildlife health. This shift in pathogen shedding with gut diversity for urban ibis may also have important consequences for the role of urban wildlife as carriers of zoonotic pathogens across the landscape. American white ibis, as well as many other wildlife species, can act as asymptomatic carriers of *Salmonella enterica* and ibis shed *Salmonella* serotypes shared with local human cases [23]. As such, a greater understanding of how wildlife microbiome form and function are altered in changing environments may be crucial for improving conservation and public health.

## Data Availability Statement

Raw sequences will be deposited at the NCBI short-read archive (https://www.ncbi.nlm.nih.gov/sra, DEPOSIT CODE TBD).

## Acknowledgements

The authors thank the University of Wisconsin Biotechnology Center DNA Sequencing Facility for providing microbial community DNA extraction and Illumina sequencing services. The authors also thank Richard Lankau for assistance with bioinformatic processing of microbiome sequences and our collaborators Sonia Altizer, Richard Hall, Jeff Hepinstall-Cymerman, Kristen Navara and Michael J. Yabsley as well as volunteers and additional undergraduate researchers for their help with study design and data collection.

## Author Contributions

### Competing Interests Statement

The authors declare no competing interests.

## Supplemental Information

**Table S1:** Landcover surrounding capture sites

|  | Number of<br>Samples | Percent Surrounding Landcover |  |  |
| --- | --- | --- | --- | --- |
|  |  | Wetland | Urban | Other* |
| JW Corbett | 1 | 91.8% | 0.0% | 8.2% |
| TetraTech | 3 | 91.8% | 0.0% | 8.2% |
| Loxahatchee Slough | 3 | 74.4% | 15.4% | 10.2% |
| Loxahatchee Wildlife Refuge | 3 | 73.8% | 4.3% | 21.9% |
| Green Cay | 5 | 26.8% | 65.5% | 7.7% |
| Kitching Creek | 2 | 15.2% | 25.5% | 59.3% |
| Solid Waste Authority | 4 | 11.1% | 49.4% | 39.4% |
| Busch Wildlife Sanctuary | 1 | 9.3% | 62.5% | 28.2% |
| Lion Country Safari | 14 | 8.9% | 46.8% | 44.3% |
| Juno Beach | 10 | 3.8% | 61.4% | 34.8% |
| Dreher Park | 9 | 0.0% | 89.7% | 10.3% |
| Dubois Park | 5 | 0.0% | 53.0% | 47.0% |
| Gaines Park | 6 | 0.0% | 84.0% | 16.0% |
| Indian Creek Park | 12 | 0.0% | 91.1% | 8.9% |
| Lake Worth | 3 | 0.0% | 76.4% | 23.6% |
| CROW Sanibel Island | 1 | - | - | - |

\*Other land classifications included agricultural, coastal, other water, and other terrestrial

**Table S2.** Summary of sample sizes and sequence outputs

| Site | Number of<br>Samples | Sequences |  | Genera |  |
| --- | --- | --- | --- | --- | --- |
|  |  | Mean<br>Number | Standard<br>Error | Mean<br>Number | Standard<br>Error |
| Busch Wildlife Sanctuary | 1 | 67605 | - | 633.0 | - |
| CROW Sanibel Island | 1 | 63960 | - | 581.0 | - |
| Dreher Park | 9 | 108408 | 15334 | 317.3 | 68.4 |
| Dubois Park | 5 | 77400 | 18890 | 315.2 | 80.0 |
| Gaines Park | 6 | 121111 | 21644 | 329.3 | 59.6 |
| Green Cay | 5 | 113303 | 32279 | 289.8 | 66.2 |
| Indian Creek Park | 12 | 115061 | 18700 | 343.4 | 52.6 |
| Juno Beach | 10 | 120359 | 11575 | 387.6 | 54.0 |
| JW Corbett | 1 | 64762 | - | 370.0 | - |
| Kitching Creek | 2 | 175378 | 62761 | 457.5 | 5.5 |
| Lake Worth | 3 | 74650 | 22398 | 351.3 | 137.4 |
| Lion Country Safari | 14 | 102983 | 18523 | 302.3 | 42.0 |
| Loxahatchee Slough | 3 | 89203 | 14894 | 410.0 | 119.1 |
| Loxahatchee Wildlife Refuge | 3 | 131894 | 31528 | 420.0 | 61.2 |
| Solid Waste Authority | 4 | 56060 | 17078 | 327.8 | 117.6 |
| Tetra Tech | 3 | 81505 | 194 | 465.0 | 27.5 |
| **All Samples** | **82** | **104695** | **6077** | **351.6** | **18.8** |

**Table S3.** Individual and site average relative abundance of dominant phyla and correlations to landscape urbanization

| Phyla | Percentage of Sequences |  |  |  | Correlation coefficient | p-value |
| --- | --- | --- | --- | --- | --- | --- |
|  | Mean (SEM)<br>(82 samples) | Mean (SEM)<br>(15 sites) | Median<br>(15 sites) | Range<br>(15 sites) |  |  |
| **Firmicutes** | **32.5 (3.1)** | **39.9 (5.3)** | **40.9** | **10.9-71.4** | **-0.773** | **<0.0001\*** |
| **Proteobacteria** | **22.6 (2.6)** | **20.2 (1.7)** | **19.0** | **12.6-36.1** | **0.772** | **<0.0001\*** |
| **TM7** | **22.6 (3.2)** | **15.2 (3.8)** | **16.7** | **0.4-37.7** | **0.763** | **<0.0001\*** |
| Actinobacteria | 6.4 (1.0) | 6.9 (1.0) | 7.5 | 2.5-14.2 | 0.131 | 0.671 |
| Chlamydiae | 6.2 (1.6) | 6.0 (2.4) | 0.8 | 0.1-34.0 | 0.297 | 0.086 |
| Fusobacteria | 3.0 (1.2) | 3.0 (0.8) | 2.0 | 0.0-8.7 | -0.261 | 0.181 |
| **Cyanobacteria** | **1.6 (0.4)** | **2.9 (1.0)** | **1.4** | **0.1-11.2** | **-0.578** | **0.004\*** |
| Chloroflexi | 1.1 (0.2) | 1.5 (0.4) | 1.1 | 0.1-4.8 | -0.514 | 0.112 |
| Tenericutes | 0.8 (0.4) | 1.4 (1.1) | 0.2 | 0.1-16.9 | -0.466 | 0.104 |
| Plantomycetes | 0.8 (0.2) | 0.9 (0.2) | 0.7 | 0.2-2.6 | -0.225 | 0.507 |
| **Bacteroidetes** | **1.1 (0.2)** | **0.9 (0.2)** | **0.9** | **0.1-2.6** | **0.528** | **0.010\*** |
| **OP11** | **0.4 (0.1)** | **0.3 (0.1)** | **0.2** | **0.0-1.1** | **0.516** | **0.007\*** |
| **TM6** | **0.3 (0.2)** | **0.2 (0.1)** | **0.1** | **0.0-1.5** | **0.457** | **0.020\*** |
| Verrucomicrobia | 0.1 (0.0) | 0.2 (0.0) | 0.1 | 0.0-0.5 | -0.067 | 0.811 |
| Acidobacteria | 0.12 (0.03) | 0.1 (0.0) | 0.1 | 0.0-0.5 | 0.037 | 0.935 |

\* Significant correlation after adjustment for false-discovery rate (Benjamini, Yoav; Hochberg, Yosef (1995). Controlling the false discovery rate: a practical and powerful approach to multiple testing. Journal of the Royal Statistical Society, Series B. 57 (1): 289–300.)

**Table S4.** Path analysis statistical results

| Regression | Estimate | SEM | p-value |
| --- | --- | --- | --- |
| Salmonella Prevalence~ |  |  |  |
| Mean Bacterial Diversity | -0.413 | 0.171 | 0.016 |
| Mean Bacterial Diversity~ |  |  |  |
| Bacterial Composition (PCoA1) | -0.392 | 0.149 | 0.009 |
| Percent Adult | 0.647 | 0.138 | <0.001 |
| Bacterial Composition (PCoA1)~ |  |  |  |
| Percent Urban Habitat | 1.180 | 0.209 | <0.001 |
| Mean Percent of Diet Provisioned | 1.514 | 0.694 | 0.029 |
| Mean Percent of Diet Provisioned~ |  |  |  |
| Percent Urban Habitat | 0.210 | 0.053 | <0.001 |
| Model fit: Overall model fit p=0.223, Comparative fit index =0.950, Tucker-Lewis index=0.900 |  |  |  |

**Figure S1.**
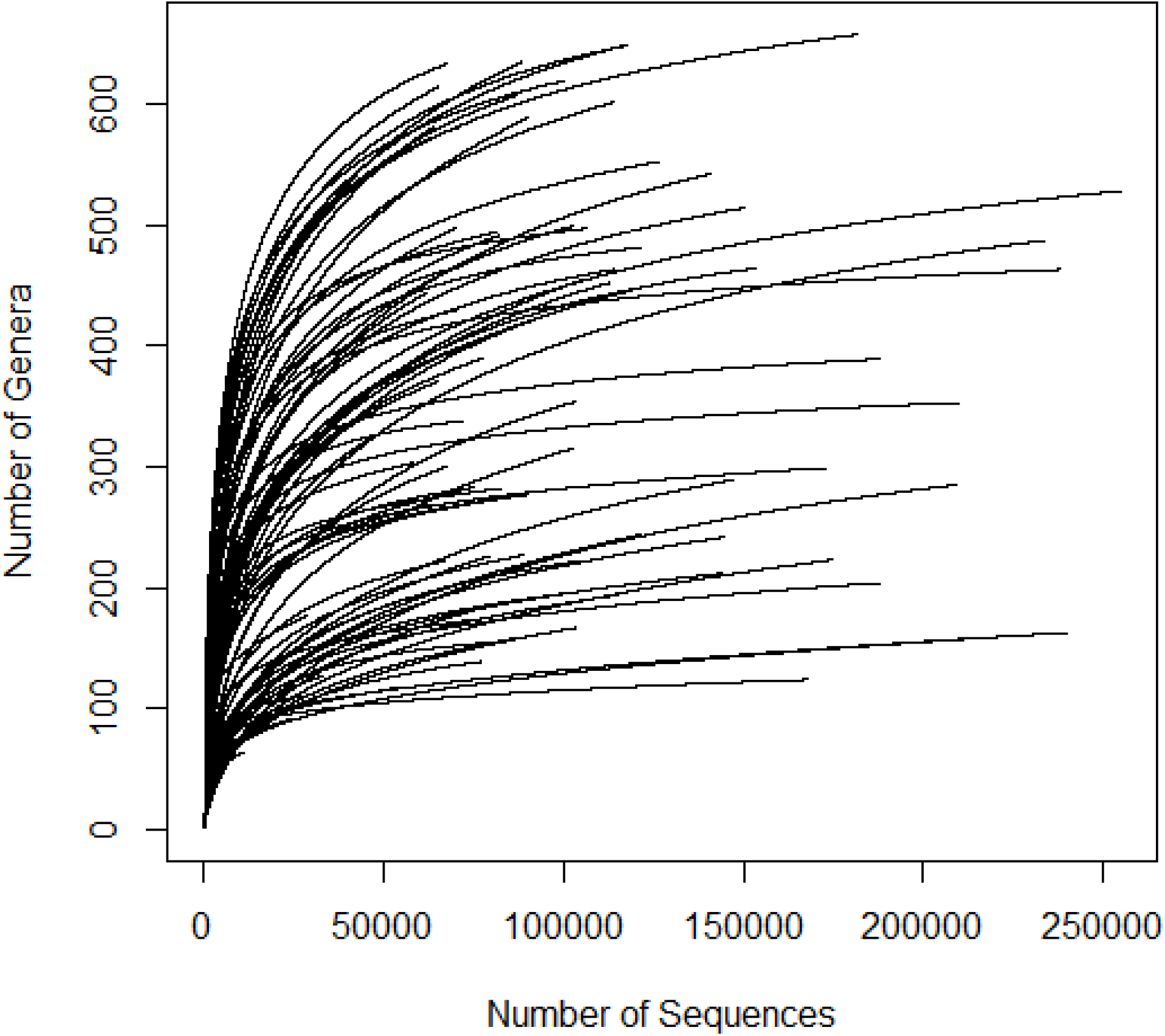
Rarefaction curves of bacterial genera detected in American white ibis feces. Rarefaction was performed using the rarecurve function in the vegan package [39].

